# Low-glutathione mutants are impaired in growth but do not show an increased sensitivity to moderate water deficit

**DOI:** 10.1101/710962

**Authors:** Sajid A.K. Bangash, David Solbach, Marcus Jansen, Fabio Fiorani, Markus Schwarzländer, Stanislav Kopriva, Andreas J. Meyer

## Abstract

Glutathione is considered a key metabolite for stress defense and elevated levels have frequently been proposed to positively influence stress tolerance. To investigate whether glutathione affects plant performance and the drought tolerance of plants, wild-type Arabidopsis plants and an allelic series of five mutants (*rax1, pad2, cad2, nrc1*, and *zir1*) with reduced glutathione contents between 21 and 63 % compared to wild-type glutathione content were phenotypically characterized for their shoot growth under control and water-limiting conditions using a shoot phenotyping platform. Under non-stress conditions the *zir1* mutant with only 21 % glutathione showed a pronounced dwarf phenotype. All other mutants with intermediate glutathione contents up to 62 % in contrast showed consistently slightly smaller shoots than the wild-type. Moderate drought stress imposed through water withdrawal until shoot growth ceased showed that wild-type plants and all mutants responded similarly in terms of chlorophyll fluorescence and growth retardation. These results lead to the conclusion that glutathione is important for general plant performance but that the glutathione content does not affect tolerance to moderate drought conditions typically experienced by crops in the field.

## Introduction

Crop yield is severely constrained by environmental stress factors resulting in a gap between the yield potential and the actual yield. The yield gap is predicted to increase in the future due to climate change and due to increasing temperature and extended phases of moderate to severe drought in particular [1-4]. Understanding the tolerance and protection mechanisms of plants is mandatory to breed crops that are able to ensure high yield even under intermittent phases of stress. Plant growth and stress defense are controlled by a multitude of different factors building tight regulatory networks that provide the plasticity that is required to ensure survival and yield of plants even under adverse conditions. The tripeptide glutathione (GSH) is considered a key metabolite in plant defense reactions against biotic and abiotic stress factors [5]. One important function of GSH is the detoxification of reactive oxygen species and organic peroxides that are frequently formed in access in stress situations [6,7]. H_2_O_2_ is in part detoxified via the glutathione-ascorbate cycle in which ascorbate peroxidases reduce H_2_O_2_ at the expense of ascorbate [8]. Dehydroascorbate ultimately resulting from ascorbate oxidation is subsequently reduced by GSH resulting in the formation of glutathione disulfide (GSSG), which is then again reduced by glutathione-disulfide reductases (GRs). The glutathione-ascorbate cycle and dehydroascorbate reductase in particular was recently shown to play a central role in minimizing drought-induced grain yield loss in rice [9]. Arabidopsis mutants lacking GR in the cytosol have been shown to have a slightly less negative glutathione redox potential (*E*_GSH_) [10]. It has also been shown that cytosolic GR plays a crucial role in leaf responses to intracellular H_2_O_2_ and in regulation of gene expression through salicylic acid and jasmonic acid signaling pathways [11].

Oxidation of two GSH molecules results in one GSSG molecule, which makes *E*_GSH_ dependent on both the degree of oxidation and the total amount of glutathione [12]. The glutathione pool in the cytosol, plastids, mitochondria and peroxisomes is, however, in a reduced state with almost 100 % GSH (in the low mM range) and only nanomolar concentrations of GSSG [13,14]. GSH is synthesized in two enzyme-dependent steps catalyzed by glutamate-cysteine ligase (GSH1) and glutathione synthase (GSH2). The biosynthetic pathway is controlled by negative feedback control of GSH on the first enzyme GSH1 [15]. With GSH as a key metabolite in stress defense reactions it is not surprising that several independent genetic screens for mutants sensitive to abiotic or biotic stresses resulted in isolation of mutants with diminished GSH content caused by mutations in GSH1 [16-21]. Mutants with intermediate GSH levels of 20 to 40 % are frequently described with a wild-type like phenotype under control conditions [18], although some reports also indicate slightly retarded growth [22]. A causal link between GSH and plant growth is particularly emphasized by the Arabidopsis mutants *zinc tolerance induced by iron 1* (*zir1*), which develops as a dwarf and contains only 15 % of wild-type GSH, and *root meristemless 1* (*rml1*) in which postembryonic development is largely abolished due to an almost complete lack of GSH [20,21].

As a key metabolite in the glutathione-ascorbate cycle and a putative cofactor for other ROS scavenging systems like glutathione S-transferases (GSTs), glutathione has frequently been linked to general stress responses in plants [5]. During drought stress responses guard cells are known to produce H_2_O_2_ that may be exploited for signaling but also needs to be detoxified to avoid serious damage [23,24]. Large-scale changes of cellular redox homeostasis and particularly of *E*_GSH_ have been considered to link primary stress responses to downstream targets [12,25]. Stress-dependent changes in *E*_GSH_ can be visualized in live cells with redox-sensitive GFP (roGFP) [13,26,27] and the question has been raised whether severe water stress might trigger an oxidative response that might be involved in drought signaling. While harsh hypo-osmotic treatments (1 M mannitol) did not cause any acute effects as measured by the roGFP2 biosensor [28]. Jubany-Mari and colleagues reported a gradual drought-induced oxidative shift of about 10 mV in the cytosol over a period of several days [29].

The latter finding raised the question whether changes in *E*_GSH_ participate in the water deficit response and whether mutants with altered *E*_GSH_ are impaired in their response. First experiments to answer this question have provided contradicting results. Overexpression of the first and regulatory enzyme in GSH biosynthesis, GSH1, in tobacco has been reported to confer tolerance to drought stress [30]. Consistent with this an increased GSH level was reported to confer tolerance to drought and salt stress while the partially GSH-deficient mutant *pad2* displays a significantly lower survival rate than wild-type plants after a two-week drought treatment [31]. In contrast, it has been reported recently that GSH deficiency in *pad2* plants does not affect the water deficit response during a 9-day drought period [32]. Similarly, it has also been reported that partial depletion of GSH in the *gsh1* mutant alleles *cad2, pad2*, and *rax1*, did not adversely affect the leaf area of seedlings exposed to short-term abiotic stress [22]. Surprisingly, the negative effects of long-term exposure to oxidative stress and high salt concentrations on leaf area were less marked in the GSH synthesis mutants than in wild-type plants. The apparent contradiction between these studies may result from different stress treatments and different scoring systems recording either survival [31] or biomass increase and leaf area [22,32]. Furthermore, the informative value of multiple studies on the role of GSH in stress tolerance is limited by the fact that frequently only individual mutants are compared to wild-type plants. Schnaubelt *et al.* considered this point by testing three different mutants, which, however, all contained intermediate levels of GSH with little phenotypic variation under non-stress conditions [22,33]. In addition, the severe stress regimes used in this study with high salt (75 mM NaCl) and osmotic stress (100 mM sorbitol) and the evaluation of clearly visible macroscopic markers is unlikely to provide a refined picture of stress sensitivity in Arabidopsis. Such a strategy can be particularly problematic when used to assess whether the growth of mutant or transgenic lines is impacted by changes in stress signaling pathways because difficulties with experimentation including possible physical damage of roots and uptake of osmotica during transfer and possible uptake and breakdown of osmotica [2,34]. Similarly, harsh drought treatment ultimately leading to wilting and death of soil-grown plants is not suitable to compare the performance of different genotypes with different growth characteristics, such as smaller plants [35].

To investigate the potential role of GSH in the response of Arabidopsis plants to moderate water deficit in more detail, we extended the allelic series of GSH-deficient mutants used by Schnaubelt *et al*. [22,33] by the latest additions, namely the mutants *nrc1* [19] and *zir1* [20] of which the latter is particularly interesting given its reported dwarf phenotype. Wild-type plants and all mutants were compared side-by-side for their growth and drought tolerance by using advanced non-invasive high-throughput shoot phenotyping enabling continuous recording of growth responses. We demonstrate that GSH-deficient mutants display diminished growth that is more severe in low GSH mutants, while even in mutants with pronounced growth deficits the decrease in GSH does not negatively impact on tolerance to moderate drought treatment.

## Materials and methods

### Plant material and growth conditions

*Arabidopsis thaliana* L. (Heyn.) ecotype Columbia-0 (Col-0) was used for all experiments as a wild-type control. Mutants with defects in GSH1 (At4g23100) were provided by the colleagues who reported their isolation and initial characterization. Arabidopsis plants were grown on a mixture of soil (www.floragard.de), sand and perlite in 10:1:1 ratio and kept in controlled growth chambers under long day conditions with 16 h light at 19 °C and 8 h dark at 17 °C. Light intensity was kept between 50 and 75 µE m^-2^ s^-1^ and relative humidity at 50 %. Seeds of wild-type and all mutants were harvested at the same time and used at a similar age for all further experiments. For initial phenotypic and physiological characterization seedlings were grown on vertically oriented agar plates under sterile conditions. Seeds were surface-sterilized with 70 % (v/v) ethanol for 5 min and plated on half-strength standard MS medium (M0222.0050; Duchefa, www.duchefa-biochemie.nl), supplemented with 0.5 % (w/v) sucrose and solidified with 0.8 % (w/v) agar. Seeds were then stratified for 2 d in the dark at 4 °C and germinated under long day conditions with 16 h light at 22 °C and 8 h dark at 18 °C. Light intensity was 75 µE m^-2^ s^-1^ and relative humidity at 50 %.

Details on growth conditions applied in large-scale non-invasive phenotyping experiments are provided in the respective method descriptions.

### Analysis of low-molecular-weight thiols

Approximately 20 mg plant material from 5-day-old seedlings grown on plates under sterile conditions was homogenized and extracted in a 10-fold volume of 0.1 N HCl. Samples were centrifuged for 10 min at 4 °C. 25 µL of the supernatant were mixed with 20 µL of 0.1 M NaOH and 1 µL of 100 mM freshly prepared dithiothreitol (DTT) to quantitatively reduce disulfides. Samples were vortexed, spun down and kept for 15 min at 37 °C in the dark. Afterwards, 10 μL 1 M Tris/HCl pH 8.0, 35 μL water and 5 μL 100 mM monobromobimane in acetonitrile (Thiolyte® MB, Calbiochem, www.merckmillipore.com) were mixed and added to the samples. The samples were vortexed, spun down and kept for 15 min at 37 °C in dark. 100 μL of 9 % (v/v) acetic acid were added, samples were vortexed and centrifuged at 13,000 g for 15 min at 4 °C. 180 µL of the supernatant were filled in HPLC vials. Thiol conjugates were separated by HPLC (SpherisorbTM ODS2, 250 × 4.6 mm, 5 µm, Waters, http://www.waters.com) using buffer C (10 % (v/v) methanol, 0.25 % (v/v) acetic acid, pH 3.7) and D (90 % (v/v) methanol, 0.25 % (v/v) acetic acid, pH 3.9). The elution protocol was employed with a linear gradient from 4 to 20 % D in C within 20 min, with the flow rate set to 1 mL/min. Bimane adducts were detected fluorimetrically (UltiMate™ 3000, Thermo Fisher, http://www.thermofisher.com) with excitation at 390 nm and emission at 480 nm.

### Phenotypic characterization of shoot growth and drought stress response

Shoot growth was analyzed automatically by using the GROWSCREEN FLUORO setup described earlier [36,37]. Seeds of WT and all *gsh1* mutant lines were stratified for 3 days at 4 °C in the dark and then placed in pots individually. Subsequently seeds were germinated and on day seven after germination seedlings with similar size were transferred into larger pots (7 × 7 × 8 cm) and randomized on trays with 30 plants on each tray. In exceptional cases plants died after this transfer and were removed from further analysis. The plants were then grown in growth chambers under fully controlled conditions at 22/18 °C, 170 µmol m^−2^ s^−1^ PAR, and 8/16 h day/night regime. The soil water content (SWC) was recorded gravimetrically. After initial soaking the SWC was allowed to decrease until a value of approx. 40 % was reached. Subsequently this SWC was held through intermittent addition of water. Starting from day 15 after sowing all plants were initially documented for the projected leaf area (PLA) and chlorophyll fluorescence every second day. During the exponential growth phase of shoots in weeks 5 and 6 after sowing all growth parameters were collected on a daily basis. All readings were taken around midday to ensure that the rosettes are oriented almost horizontally above the soil. The PLAs A_1_ and A_2_ of two consecutive days were used to calculate the relative shoot growth rate (RGR_Shoot_) (% d^-1^) according to the equation RGR_Shoot_ = 100 × 1/t × ln(A_2_ /A_1_). Chlorophyll fluorescence was recorded after dark adaptation for at least 30 min with a camera-based system to calculate color coded images of F_v_ /F_m_ as a measure of the potential quantum yield of photosystem II. For further analysis, average values for F_v_ /F_m_ for whole rosettes were calculated.

For drought stress experiments the plants were split into two subpopulations of which the first population was well-watered throughout the experiment while the second population was exposed to drought from day 24 onwards until growth ceased on day 37. Subsequently plants were again watered to a soil water content of about 40 % and allowed to recover. All plants were harvested after 44 days to determine fresh and dry weight.

### Determination of rosette morphology

Leaf circumference, rosette compactness, rosette stockiness, and eccentricity were calculated from the PLA as described earlier [36].

### Ratiometric roGFP2 imaging

5-day-old Arabidopsis seedlings stably expressing Grx1-roGFP2 in the cytosol were imaged by CLSM. Seedlings were mounted on a glass slide in a drop of distilled water and immediately transferred to a Zeiss LSM780 confocal microscope. Images of root tips were collected with a 40x lens (Zeiss C-Apochromat 40x/1.2 NA water immersion). RoGFP2 was excited with the 405 and 488 nm laser lines in multi-track mode with a pixel size of 0.415 µm in x and y and 1.58 µs pixel dwell time with line switching and averaging four scans per line. The pinhole was set to 73 µm (2.0 - 2.3 Airy units dependent on the wavelengths). Emission was collected from 504 to 530 nm. Autofluorescence excited at 405 nm was collected from 431 to 470 nm. Images were processed using a custom built MATLAB tool with x,y noise filtering, fluorescence background subtraction and autofluorescence correction as described previously [38].

### Statistical analysis

Statistical analysis was performed using the software GraphPad Prism 7. Different data sets were analyzed for statistical significance with the One- and two-way Analysis of Variance (One- and two-way ANOVA) followed by Tukey’s multiple comparisons test.

## Results

### Severe osmotic stress imposed by mannitol causes partial oxidation of cytosolic glutathione

To investigate the impact of osmotic challenge on Arabidopsis plants, we initially grew seedlings expressing the *E*_GSH_-sensor Grx1-roGFP2 in the cytosol on 300 mM mannitol to mimic severe drought stress. In addition to wild-type seedlings, the effect of mannitol on *E*_GSH_ was also studied in *gr1* mutants deficient in cytosolic GR because impaired reduction capacity for GSSG renders the glutathione pool more sensitive to stress-induced oxidation [10]. While the 405/488-nm fluorescence ratio for Grx1-roGFP2 in wild-type root tips was close to values for full reduction measured after incubation with DTT, the readout for the cytosol of *gr1* roots was significantly higher with intermediate ratios between full reduction and full oxidation of the sensor (Fig 1A). From these measurements cytosolic redox potentials of about -310 mV in the wild-type and close to the midpoint of the sensor at -280 mV for *gr1* can be deduced. In both cases, for wild-type and *gr1* seedlings, treatment with 300 mM mannitol did not cause significant changes in the fluorescence ratio during the first 16 h. This indicates that within this time window the seedlings were still fully capable of maintaining their thiol redox homeostasis in the cytosol. In contrast to this observation, continuous growth on 300 mM mannitol for 5 days caused a pronounced ratio shift in wild-type and *gr1* root tips. While the ratio values in wild-type root tips reached intermediate values, the ratios in *gr1* root tips approached full oxidation in the root cap while the meristematic region appeared still partially reduced (Fig 1B).

**Fig 1.**
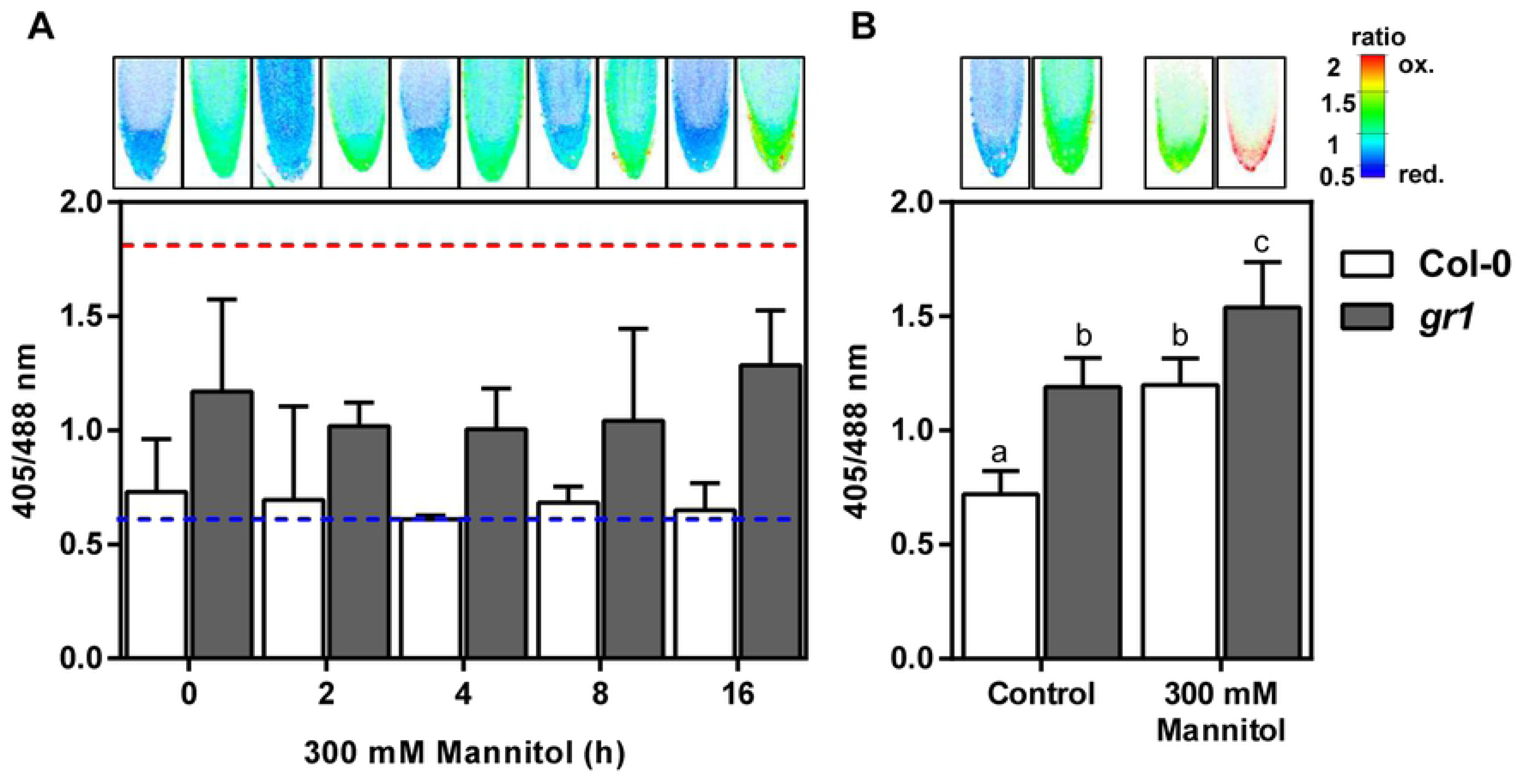
Long-term osmotic stress causes partial oxidation of cytosolic Grx1-roGFP2 in Arabidopsis root tips. (**A**) 5-day-old seedlings transferred to 300 mM mannitol. The dashed horizontal lines indicate the ratio values resulting from treatment with 10 mM DTT for full reduction (blue) and 25 mM H_2_O_2_ for full oxidation (red). (**B**) 5-day-old seedlings germinated and continuously grown on 300 mM mannitol. All values are means ± SD (*n* ≥ 10). Letters indicate significant differences (One-way ANOVA with Tukey’s multiple comparisons test; *p* ≤0.05).

Based on our observation that long-term drought stress impacts on *E*_GSH_ and causes oxidation in the cytosol, as well as on published reports on decreased drought tolerance of GSH-deficient mutants [31], we set out to test the hypothesis that the GSH content of Arabidopsis plants correlates with drought sensitivity.

### An allelic series of *gsh1* mutants

Several genetic screens have led to identification of different *gsh1* mutant alleles that have all been reported to be GSH-deficient albeit at different degrees. For the degree of GSH depletion, however, values between 20 and 40 % GSH compared to wild-type plants are frequently cited, which precludes unambiguous ranking of the mutants according to their GSH content. While GSH levels may indeed vary between growth conditions, the annotation of mutants with concentration range of GSH makes selection of the most appropriate alleles for comparative experimental work difficult. To rank all five available mutants of the allelic series that can be maintained in their homozygous state according to decreasing GSH content, seedlings were grown under controlled conditions on agar plates and analyzed for their GSH content. The HPLC analysis revealed a separation of wild-type and the different *gsh1* mutants into 3 distinct classes (Fig 2A). Wild-type seedlings contained 273 ± 39 nmol g^-1^ FW glutathione and *rax1* seedlings 142 ± 46 nmol g^-1^ FW. The third class included *pad2, cad2, nrc1* and *zir1* which all contained less than 73 nmol g^-1^ FW glutathione. Although not statistically different the measured mean content of glutathione in *zir1* was only 45 ± 20 and thus lower than in the other three mutants of this class. Depletion of GSH due to mutations in GSH1 was accompanied by a concomitant increase of cysteine as one of the GSH1 substrates (Fig 2B).

**Fig 2.**
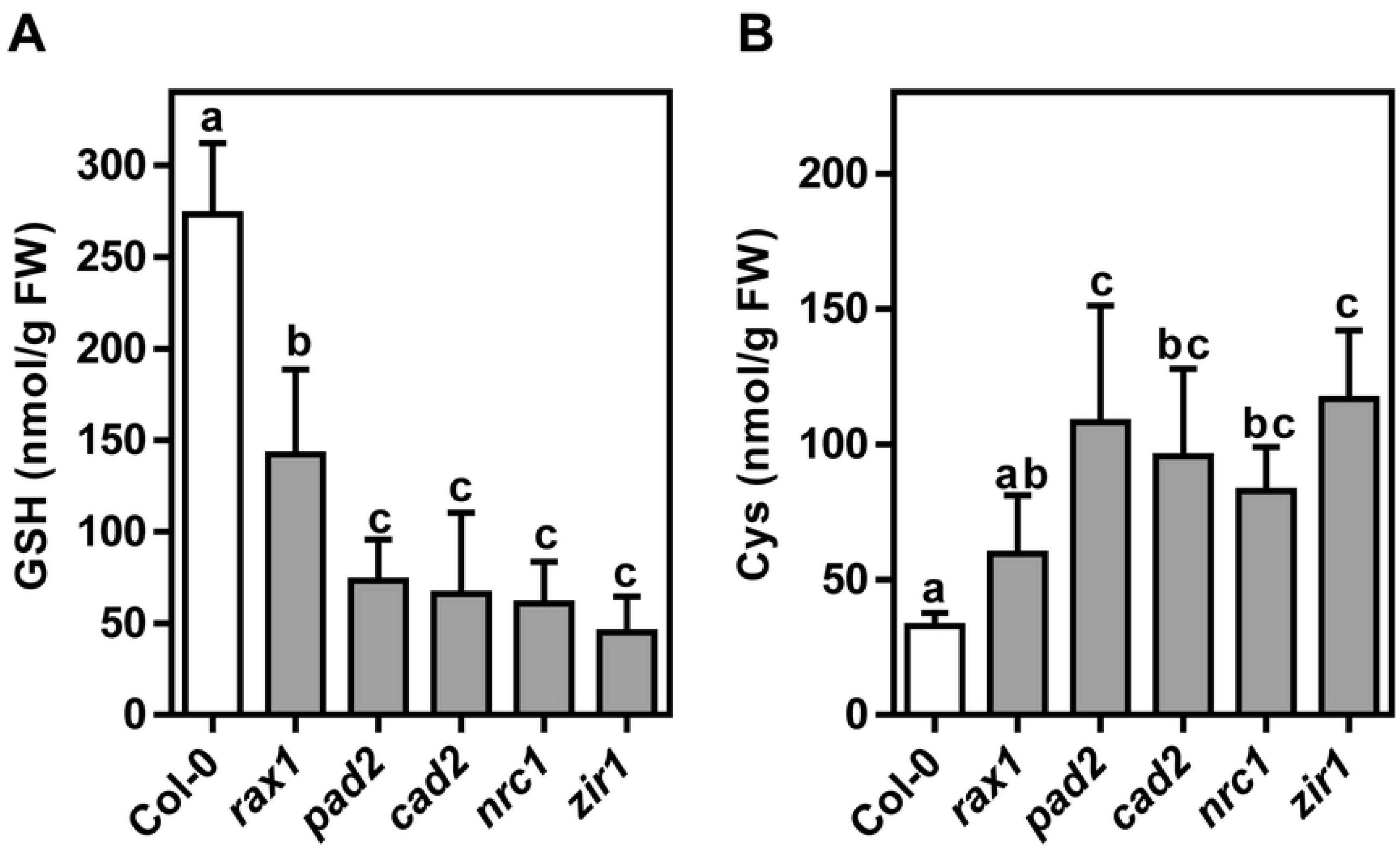
Glutathione content in a series of allelic mutants deficient in glutamate-cysteine ligase. (**A,B**) HPLC analysis of total glutathione presented as GSH equivalents (**A**) and cysteine (**B**) in 5-day-old seedlings. The presented values are means ± SD (*n* = 5). Lower case letters indicate statistically different values (One-way ANOVA with Tukey’s multiple comparisons test; p < 0.05).

### Soil-grown GSH-deficient mutants display growth phenotypes but are insensitive to moderate water deficit

To study the relationship between glutathione content and biomass gain under drought, we analyzed the growth of *gsh1* mutants and wild-type in a sub-lethal drought stress assay. A large population of soil-grown plants was separated into two sub-populations of which one was used as a well-watered control group while the second was exposed to drought stress until growth ceased (Figs 3A, 4 and 6B). At the end of the drought period plants did not yet show obvious signs of wilting (Fig 4). After the drought period, all plants were watered again reaching a SWC of about 40 - 50 % for another week before the plants were harvested (Fig 3A). Within the population of plants grown with sufficient water supply throughout the entire growth period no distinct growth phenotype could be observed for *cad2* (Figs 3B and 5C). The glutathione-deficient mutants *rax1, pad2* and *nrc1* were consistently smaller than wild-type plants by up to 30 % at the end of the 6-week growth period (Figs 3B and 5C). Growth retardation was apparent already on day 22 after sowing (Fig 5A). In contrast to mutants with intermediate levels of GSH, *zir1* showed a distinct dwarf phenotype with only 19 % of the wild-type biomass.

**Fig 3.**
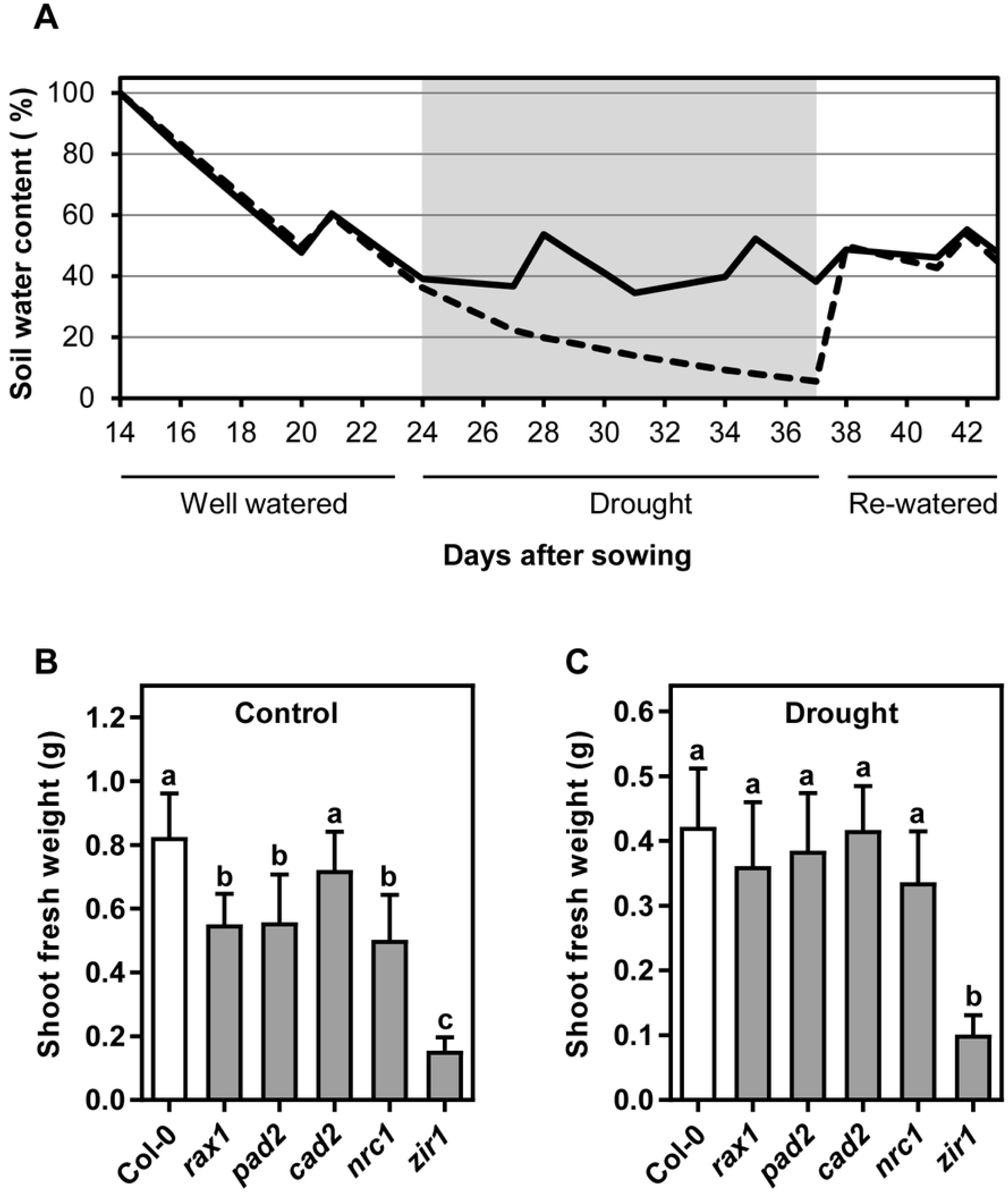
Drought-induced growth reduction is not affected by the GSH content of Arabidopsis plants. **(A**) Watering regime and soil water content during the experiment for drought stressed plants (dashed line) and control plants (solid line). (**B**,**C**) Shoot fresh weight under control and drought conditions respectively at the end of the growth period after 6 weeks. Plants were grown in soil-filled pots under short day conditions and exposed to water stress as illustrated in panel **A**. Values are means ± SD (*n* ≥ 10). Letters in each graph indicate significant differences (One-way ANOVA with Tukey’s multiple comparisons test; *p* < 0.05).

**Fig 4.**
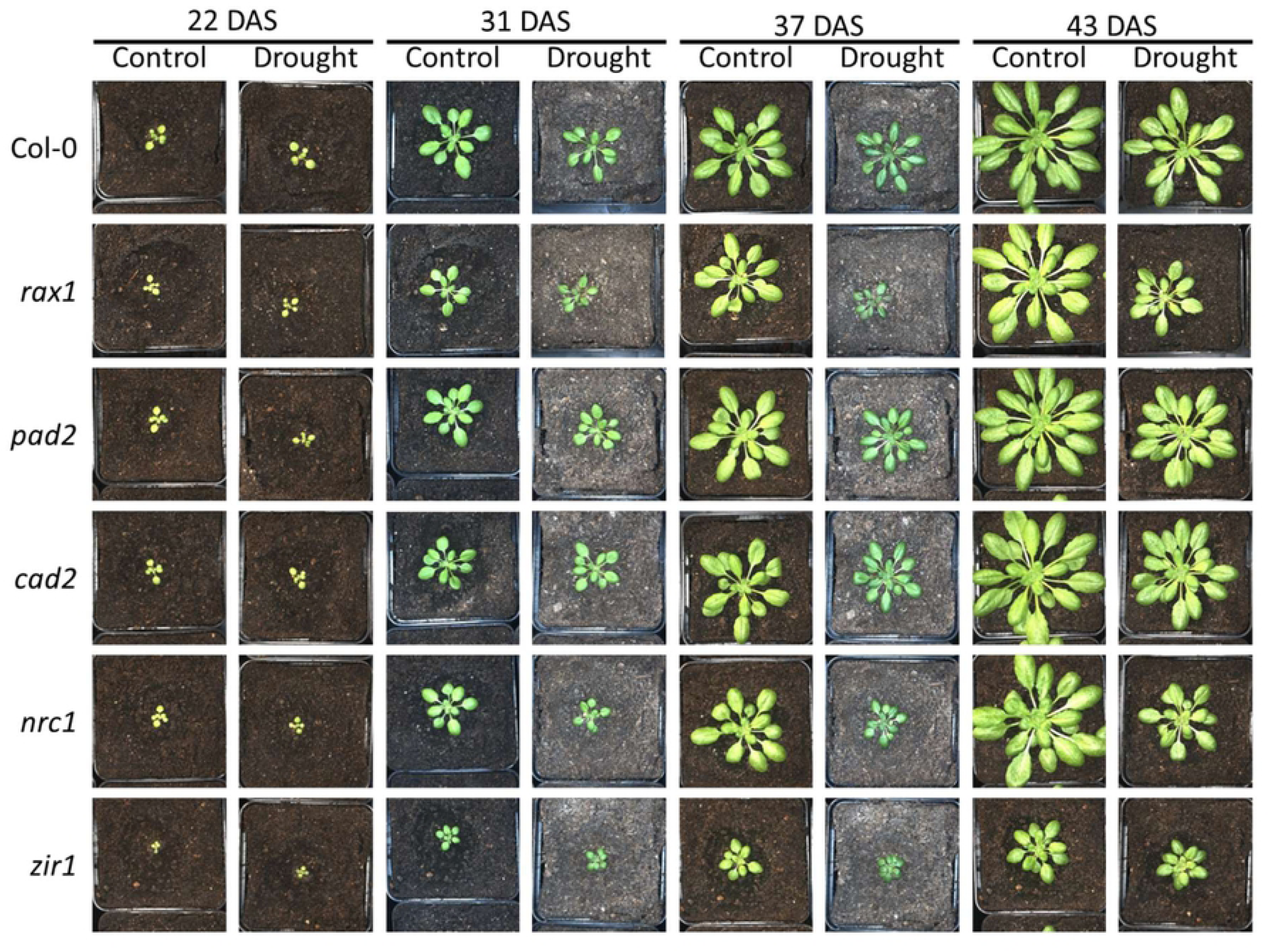
Photographs of representative plants during the drought stress experiment. DAS: days after sowing.

**Fig 5.**
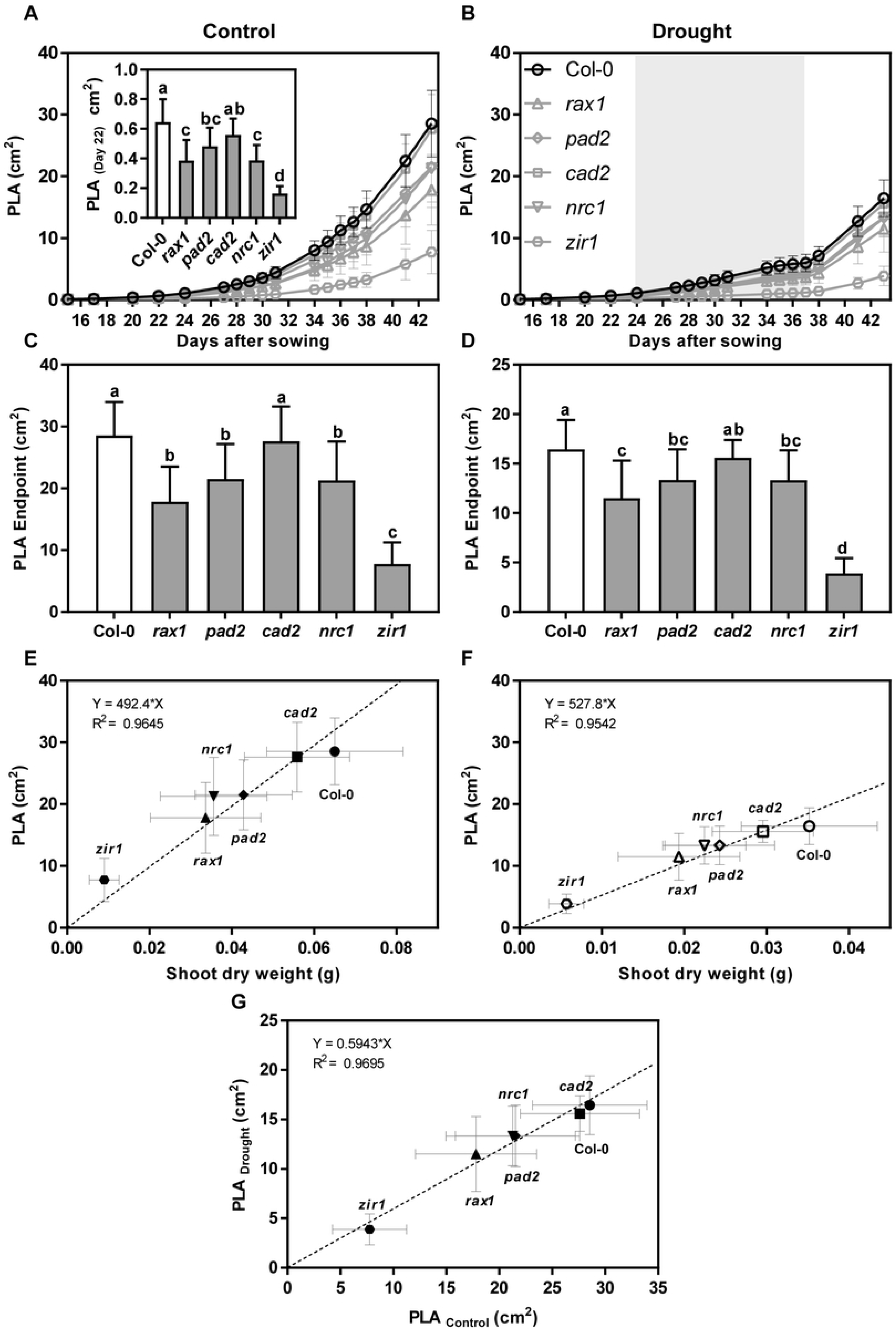
Projected leaf area (PLA) in wild-type and *gsh1* mutants grown under control and drought stress conditions in soil-filled pots. (**A**,**B**) Continuous recording of PLA during the entire growth period for well-watered control plants (**A**) and drought-stressed plants (**B**). The inset in (**A**) shows the PLA of well-watered plants 22 DAS to emphasize that different growth could be seen here already. Identifiers for the different growth curves are provided in (**B**). (**C**,**D**) PLA at the time of harvest 44 DAS for control (**C**) and drought-stressed (**D**) plants. (**E**,**F**) PLA for wild-type and all *gsh1* mutants under control (**E**) and water deficit conditions (**F**) measured at the end of the growth period. For calculation of the regression the origin of co-ordinates (point 0/0) was included as an additional virtual data point. (**G**) Relative PLA. The calculated linear regression indicates a direct correlation between PLA under drought and control conditions for all plant lines under investigation. All values are means ± SD (*n* ≥ 16 randomly distributed on 8 trays).

Lack of water supply for an interim period of 13 days during the entire growth phase of 44 days caused a pronounced growth retardation in wild-type plants by about 50 % as compared to the well-watered controls recorded as a decrease in shoot fresh and dry weight as well as PLA (Figs 3B, 3C, 5A-D and S1A and S1C Figs). The shoot water content at the end of the experiment was similar in control and drought-stressed plants indicating that the applied drought stress did not cause serious damage, such as permanent lesions (S1B and S1D Figs). Direct comparison of drought-stressed *gsh1* mutants and the respective wild-type revealed that for the intermediate GSH-deficient mutants the relative retardation was less pronounced than under control conditions or could not even be detected statistically (Figs 3C, 5D and S1C Fig). The major exception with the most pronounced difference to the wild-type is *zir1* for which the retardation as clearly visible in all recorded parameters. The severe dwarf phenotype of *zir1* enabled further analysis of the PLA at the end of the experiment for all mutants despite the lack of consistent phenotypic differences for the intermediate mutants. Comparison of the PLA with the dry weight (DW) for well-watered control plants and drought-stressed plants resulted in linear relationships indicating that the specific leaf area (PLA g^-1^ DW) in each group of plants is not affected by glutathione deficiency (Figs 5E and 5F). To further test whether mutants with very low GSH are more seriously affected by drought than mutants with intermediate levels of GSH and wild-type plants, we plotted the PLA of drought-stressed plants against the PLA of control plants. The linear relationship strongly indicates that even low GSH mutants that grow as dwarfs are not severely affected in their ability to withstand moderate water deficit (Fig 5G).

Drought-induced growth retardation was also apparent from the continuous recording of the PLA (Figs 5A-D) and the RGR_Shoot_ calculated from these measurements (Figs 6A and 6B). Regular measurement of the PLA over the entire growth period also enabled the calculation of the relative shoot growth rate (RGR_Shoot_) on a daily basis. RGR_Shoot_ in the control population gradually decreased from 28 % d^-1^ at the start of the recordings on day 16 after sowing to 12 % d^-1^ at the end of the growth period (Fig 6A). Over the entire growth period no obvious deviations in RGR_Shoot_ were apparent between the wild-type and the different *gsh1* mutants with intermediated GSH content. For *zir1* RGR_Shoot_ showed a similar decrease over time, but there were a few intermittent days with significantly lower RGR_Shoot_ than in all other plants (Fig 6A). In the drought-stressed population RGR_Shoot_ decreased significantly faster during the growth period approaching zero after 13 days of water withdrawal (Fig 6B and 6C). The drought stress period started by chance during a phase when *zir1* already showed particularly low values for RGR_Shoot_ in the control population (Fig 6A and 6B). From this point onwards RGR_Shoot_ was consistently lower in *zir1* compared to all other lines for almost the entire drought period (Fig 6B). Immediately after re-watering on day 37 the RGR_Shoot_ increased to peak values of about 20 % d^-1^ on day 41 and subsequent decline to values of about 15 % d^-1^ on day 43 for all lines including *zir1*. After re-watering, the glutathione-deficient mutants even showed a tendency of faster growth compared to wild-type plants (Fig 6B and 6C).

**Fig 6.**
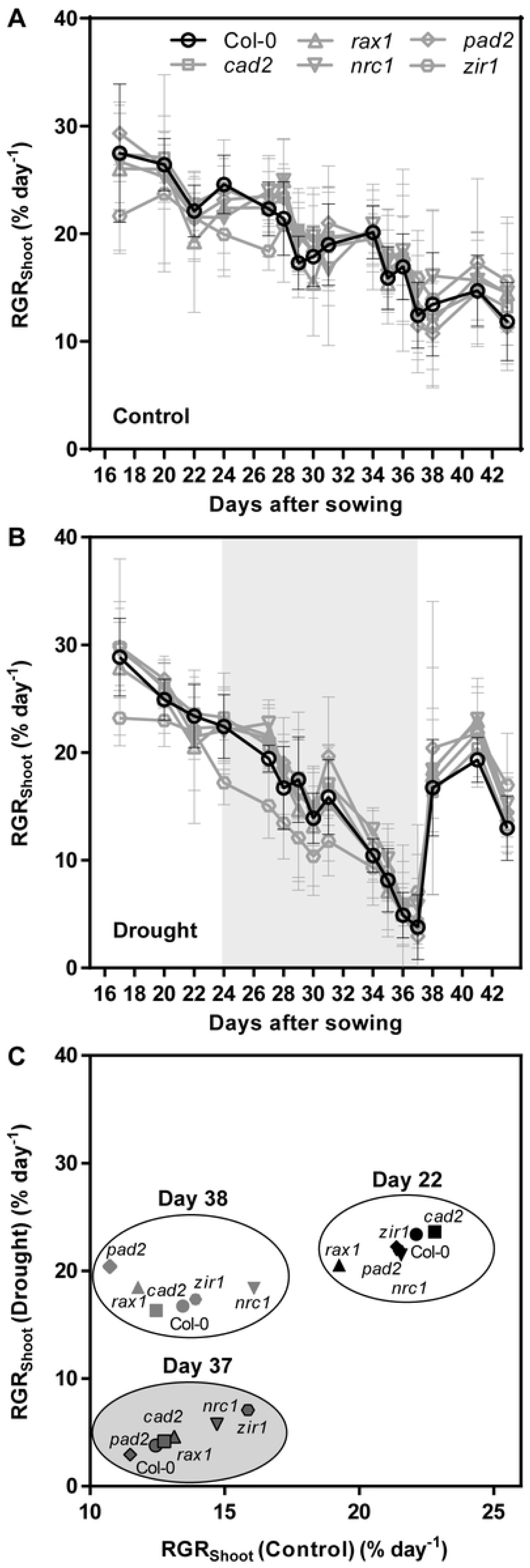
Relative shoot growth rate (RGR_Shoot_) in wild-type and *gsh1* mutants grown und control and drought-stress conditions. Continuous recording of RGR_Shoot_ during the entire growth period for control (**A**) and drought-stressed plants (**B**). The period of water withdrawal for the drought-stressed population is indicated by a grey shadow. Water withdrawal ended when RGR_Shoot_ approached zero on day 37. (**C**) Comparison of relative growth rates for plants in well-watered and drought stressed populations at three critical time points during the experiment. Symbols for the different lines are used as described in panel **A**.

With each measurement of all individual plants the potential quantum yield of photosynthesis (F_v_ /F_m_) was recorded after dark adaptation for at least 30 min. From measurements of the whole rosette single average values for the respective rosettes were calculated. In plants kept under well-watered control conditions throughout the entire growth phase, F_v_ /F_m_ reached values between 0.71 and 0.72 for wild-type plants and for the mutants with intermediate levels of GSH (*rax1, pad2, cad2* and *nrc1*). Slightly but significantly lower values for F_v_ /F_m_ of 0.70 to 0.71 were only found in the more severely GSH depleted *zir1*. While the differences between control plants and drought stressed plants were very small in all lines under investigation there was a tendency towards slightly increased values for F_v_ /F_m_ in drought-stressed plants both at the last day of the drought period and just before harvest at the end of the experiment after full recovery of drought stressed plants (Figs 7A and 7B). This difference reached significance only for *pad2, cad2* and *nrc1* on the last day of the drought phase and in *cad2* at the time of harvest.

**Fig 7.**
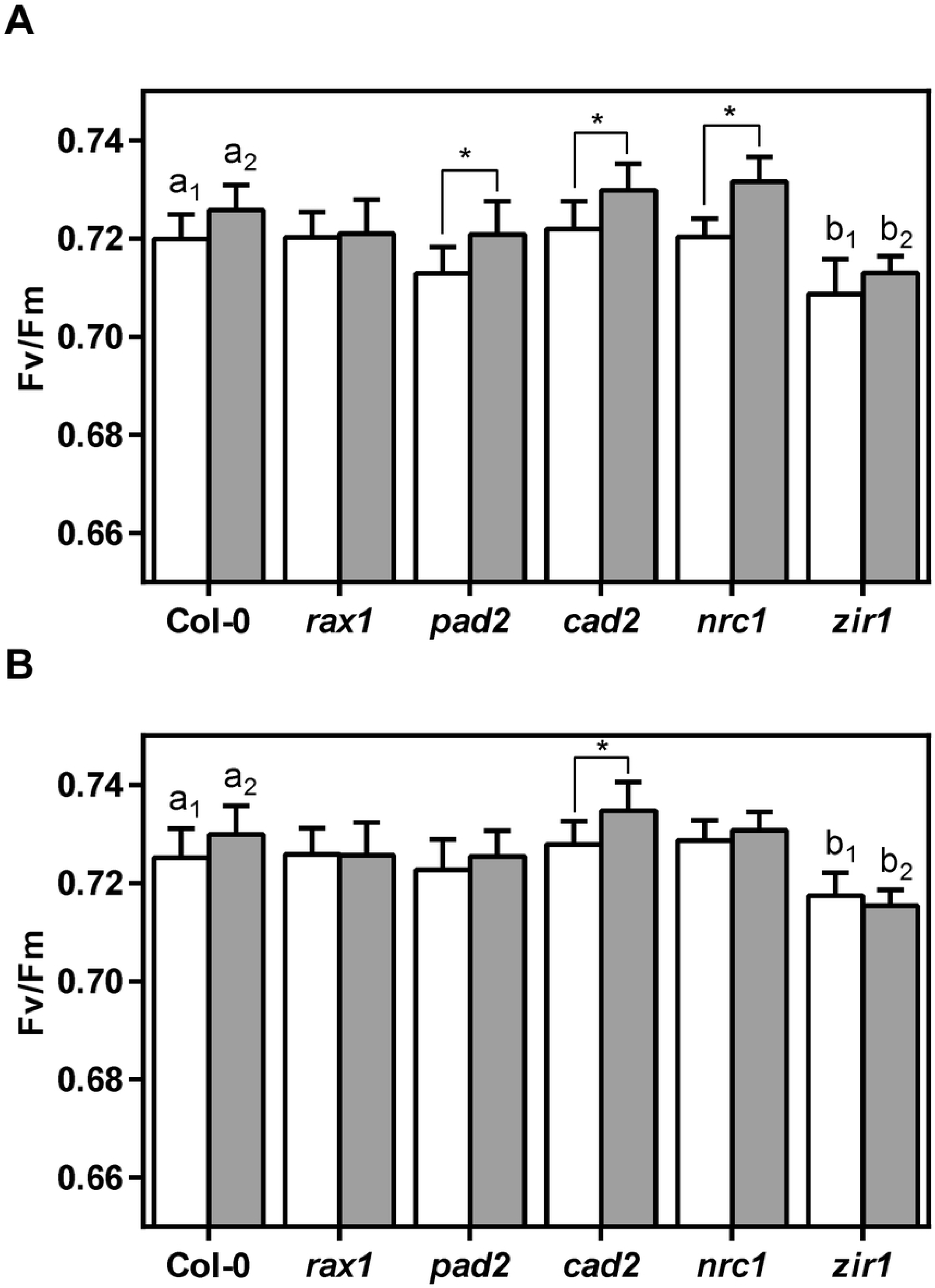
Potential quantum yield of PSII (F_v_ /F_m_) of Arabidopsis wild-type and *gsh1* mutants grown under control and drought stress conditions. (**A**) F_v_ /F_m_ on the last day of the drought period 37 DAS. (**B**) F_v_ /F_m_ at the end of the experiment 43 DAS. All values are means ± SD (*n* ≥ 16 plants). Letters in each graph indicate significant differences as determined by Two-way ANOVA with Tukey’s multiple comparisons test; *p* < 0.05). For Col-0 and *zir1* control plants and drought-treated plants were compared separately (indicated by index numbers). Values for plants grown under control conditions are shown in white and values for plants exposed to water deficit are shown in grey bars.

Recording of PLA and subsequent image analysis enabled the extraction of additional morphological characteristics of the rosette shape. These factors, included the area by circumference, the compactness of rosettes as a measure for the length of petioles, the rosette stockiness as a measure of how indented the rosette is, and the eccentricity as a parameter describing the shape of the rosette compared to a circle. Generally, all parameters reflected the measurements of PLA (S2 Fig). The only exception was *zir1* which showed an increased stockiness compared to all other lines. Due to the earlier decrease in RGR_Shoot_ *zir1* also developed a more compact rosette, a plateau in rosette stockiness and a more pronounced eccentricity after onset of drought.

## Discussion

### Glutathione affects growth under non-stress conditions

Side-by-side comparison of all available viable *gsh1* mutants for their glutathione content revealed different levels in the different mutants. The observation that *rax1* contains more glutathione than *cad2* and *pad2* confirms an earlier report by Parisy et al. [18] and the lowest glutathione content in *zir1* is in line with the original report of this mutant [20]. *nrc1* has very similar glutathione levels as *cad2* which is in line with their similar Cd^2+^-sensitivity [19]. Phenotypic comparison showed that growth of all mutants except *cad2* was impaired albeit to different degrees. *zir1* mutants were significantly smaller than all others. Comparison of our results with findings from earlier reports on individual mutants or side-by-side comparison of a subset of mutants revealed some pronounced differences. While Schnaubelt et al. [22] reported 14-day-old shoots of *rax1, pad2* and *cad2* to be significantly smaller than wild-type shoots, the original reports on the initial characterization of *cad2, rax1*, and *pad2* all found no phenotypic difference between mutant and wild-type seedlings [16-18]. While Ball et al. [17] found no effect of light on the phenotypic appearance of *rax1* at all developmental stages, a slight retardation of *pad2* seedlings observed under low light conditions got reverted under high light [22]. GSH levels may vary significantly between experiments due to some non-controlled experimental differences [18]. Thus, the use of a complete allelic series of mutants grown under exactly the same conditions in an automated phenotyping setup should avoid such limitations with lab-based phenotyping and provide more robust data. Comparison of shoot phenotypes further supports the conclusion that phenotypes are not linearly correlated with the GSH content as long as the GSH content can be kept above a certain threshold [20]. A possible explanation for the lack of a linear correlation between glutathione content and growth may be that the available mutants originate from different EMS-treated populations that were screened for sensitivities to different stress factors. This implies that, despite several rounds of backcrossing, the mutants may still contain additional cryptic mutations that are not linked to glutathione homeostasis. Such mutations may affect growth properties in a pleiotropic way similar to recent findings on ascorbate-deficient mutants [39]. In any case, this further emphasizes the added value of investigating several allelic mutants with slight deviations in GSH particularly in the low GSH range side-by-side.

The lower threshold for maintaining a wild-type-like phenotype is clearly passed in the *zir1* mutant resulting in pronounced dwarfism [20]. Glutathione is a key metabolite with essential functions in detoxification, cellular redox homeostasis and as a co-factor [5,40]. GSH is a co-factor in detoxification of methylglyoxal [41] and Fe-S cluster transfer by glutaredoxins [42]. In addition, GSH acts as a S-donor in glucosinolate biosynthesis as well as a co-substrate of ascorbate peroxidases and glutathione S-transferases (GSTs) for peroxide detoxification and of GSTs for conjugation of electrophilic xenobiotics [43-46]. Beyond this glutathione is the most abundant low molecular thiol with low millimolar concentrations in the cell [47] and as such together with glutathione reductases important for maintaining highly reducing redox potentials in the cytosol, plastids, mitochondria and peroxisomes [10,13,14]. With this multitude of functions, it is not surprising that a decrease in GSH levels eventually impair some processes, even though this may be indirectly. Although the experiments reported here were not designed to answer questions on which molecular processes are impaired, the results nevertheless indicate that a certain threshold of GSH depletion needs to be reached before growth is impaired. Below this threshold the developmental phenotype is strongly dependent on the GSH concentrations as evidenced by the *zir1* mutant but also the *rml1* mutant, which has less than 5 % GSH and arrests in growth after germination [21]. The availability of at least one viable mutant with low GSH, which shows already strong and consistent growth impairment, appears particularly useful to further test hypotheses regarding a causal link between GSH content and stress sensitivity.

### Glutathione content and drought tolerance

Plants reprogram their metabolism and growth when exposed to water limitation [2] and these changes in growth and physiological status can often be monitored non-invasively at the level of whole plants [48]. The potential quantum yield of photosynthesis F_v_ /F_m_ is a parameter for the functional status of photosystem II and it is well known that severe drought stress causes a decline in F_v_ /F_m_ [49,50]. At the same time, it is also established that F_v_ /F_m_ during the growth process increases with leaf size due to a relative increase of the lamina which typically shows high F_v_ /F_m_ values compared to leaf borders with low F_v_ /F_m_ value [36]. The slightly increased F_v_ /F_m_ in plants exposed to moderate water deficit might thus correspond to reduced growth and concomitant earlier leaf differentiation and hence a higher fraction of leaves with F_v_ /F_m_ values typical for full-grown leaves [36]. Similarly, the lower F_v_ /F_m_ values for *zir1* mutants at the end of the drought period and still after a recovery phase may thus be a consequence of delayed development and less matured leaves compared to wild-type and all other mutants. Alternatively, the lower F_v_ /F_m_ with values around 0.70-0.71 may also be indicative of partial photoinhibition in *zir1*. Photoinhibition and concomitant production of reactive oxygen species (ROS) is a common feature of stress responses. Given that GSH is involved in detoxification of ROS via the glutathione-ascorbate cycle the growth phenotype appears to be consistent with reduced growth in the ascorbate-deficient mutant *vtc2-1* [51]. This apparent link between ascorbate content and plant growth, however, has been questioned recently after identification and characterization of a true null allele *vtc2-4* that despite low ascorbate content has a wild-type-like phenotype [39].

External supply of 400 µM GSH has been reported to render Arabidopsis plants more drought tolerant [52]. Vice versa *pad2* mutants are reported to be more sensitive to drought than wild-type plants [31]. Our results, however, show no such drought sensitivity. Instead, our systematic approach rather indicates that soil-grown Arabidopsis plants exposed to moderate water deficit generally respond with a gradual decline in growth but no signs of wilting at the point of complete growth arrest. This response is not affected by the GSH content. This is even true when GSH is down to 21 % of wild-type levels, which causes severe growth retardation in *zir1* under non-stress conditions. Direct comparison of wild-type plants and mutants with GSH levels of 62 to 21 % compared to wild-type all have similar specific leaf areas, i.e. PLA/DW ratios, irrespective of whether plants had been grown under control conditions or exposed to drought stress (Fig. 5). A temporary phase of moderate water deficit caused by lack of water supply for 13 days led to a pronounced decrease in growth, but even a slight increase in the specific leaf area. This result indicates that plants that experienced temporary water deficit subsequently grew slightly faster than well-watered control plants. Re-growth after rewatering has been accounted for by cell expansion resulting in actual growth and not only a reversible change due to increased turgor [53]. Interestingly, plants exposed to mild drought stress have been observed to accumulate sugars and starch [54]. This may contribute to a quick restart of growth after re-watering in all lines including *zir1*. Important in this context is the observation that the linearity for the specific leaf area across all mutants and the wild-type is maintained, which indicates that even the very low glutathione mutant *zir1* did not get severely damaged during the stress treatment.

## Conclusion

Our analysis provides a systematic, quantitative foundation to the observation that glutathione deficiency causes retarded growth of both roots and shoots. Direct side-by-side comparison of mutants with different degrees of glutathione depletion show no gradual decrease in growth, but a minor retardation, which is similar for all mutants with 25-62 % GSH and more severe only for mutants with only 21 % GSH. This suggests that under non-stress conditions partial depletion of the cellular glutathione pool is tolerable while passing a threshold below about 25 % GSH leads to gradual impairment of plant growth.

Our systematic analysis did not show any indication that GSH content of Arabidopsis plants correlates with drought resistance. As such it contrasts earlier reports. Moderate water deficit applied through water withdrawal until shoot growth ceased showed that wild-type plants and all mutants responded similarly in terms of all morphological parameters analyzed, as well as the photosynthetic machinery as analyzed by chlorophyll fluorescence. Taken together the results indicate that glutathione is important for general plant performance, but does not affect tolerance to moderate drought conditions typically experienced by crops in the field.

## Acknowledgements

This work was supported by Deutsche Forschungsgemeinschaft (DFG) within the Research Training Group GRK 2064 (AM.; MS), grant ME1567/9-1/2 within the Priority Program SPP1710 (AM), the Emmy-Noether programme (SCHW1719/1-1; MS), and Excellence Initiative (EXC 1028; SK). The Seed Fund grant CoSens from the Bioeconomy Science Center, NRW (AM; MS) is gratefully acknowledged. The scientific activities of the Bioeconomy Science Center are financially supported by the Ministry of Innovation, Science and Research within the framework of the NRW Strategieprojekt BioSC (No. 313/323-400-002 13). SB received financial support through a fellowship from the Higher Education Commission, Pakistan. We thank Bastian Welter (University of Cologne) for technical assistance with HPLC measurements, Silvia Braun and Thorsten Brehm (FZ Jülich) for support of the shoot phenotyping and drought stress experiments, and Dr. Kerstin Nagel for discussion of phenotyping approaches and critical reading of the manuscript.

## Supporting information captions

**S1 Fig. Shoot dry weight and shoot water content for wild-type and *gsh1* mutants.** (**A**,**C**) Shoot dry weight for well-watered control plants (**A**) and drought-stressed plants (**C**). (**B**,**D**) shoot water content under control (**B**) and drought conditions (**D**). Values are means ± SD (*n* ≥ 10). Letters in each graph indicate significant differences (One-way ANOVA with Tukey’s multiple comparisons test; *p* < 0.05).

**S2 Fig. Morphological characteristics of rosettes grown under control and drought-stress conditions.**

